# Sexual division of labour shapes hunter-gatherer spatial ranges

**DOI:** 10.1101/2025.04.15.649057

**Authors:** Cecilia Padilla-Iglesias, Dieudrice Nganga, Eustache Amboulou, Juliette Ruf, Suspense Averti Ifo, Lucio Vinicius, Andrea Bamberg Migliano

## Abstract

Mobility lies at the adaptive core of the hunter-gatherer foraging niche, and has shaped the cultural and genetic evolution of our species. Yet, the specific drivers and consequences of mobility are still debated. Here we analyse the lifetime mobility patterns of 776 Mbendjele BaYaka hunter-gatherers from five regions in the northern Republic of Congo, revealing pronounced gender differences in spatial behaviour. While men expand their spatial ranges from adolescence to adulthood, women’s ranges remain stable. We find evidence for a sexual division of labour underpinning these patterns, with men’s greater mobility driven by subsistence activities such as hunting or exploratory trips away from camps, and women’s mobility mostly driven by gathering closer to their residential location. Our findings challenge traditional assumptions of patrilocality, suggesting that men travel further due to their spatial separation from kin. Larger spatial ranges are associated with increased reproductive success for both genders, suggesting adaptive benefits of mobility in accessing resources, social networks, and potential mates. By linking individual behaviors to broader movement dynamics, our study deepens our understanding of gendered mobility in humans and highlights its significance to social structuring in our species and its evolutionary consequences.

## Introduction

Hunter-gatherer multi-level social structures have key properties that foster the evolution and accumulation of cultural and genetic diversity^1–4^. Such structures are believed to underpin several unique features of *Homo sapiens*, including sexual division of labour and biparental provisioning allowing multiple dependent offspring simultaneously^5–7^; multilocality (dispersal of both genders), or co-residence and cooperation between unrelated individuals^8–10^. The flexibility and fluidity that characterises such social structures are created through the movement of individuals and groups^10,11^. Thus, given that all humans lived as hunter-gatherers until the Neolithic, mobility patterns likely coevolved with the unique suite of adaptations of our species^5,12^.

Hunter-gatherer mobility has been traditionally understood through the lens of resource distribution, with residential mobility interpreted as a necessary adaptation of small-scale bands against resource depletion^13–16^. However, hunter-gatherer social networks may be far more extensive than previously assumed, encompassing broad geographical ranges and frequent interactions with a large number of individuals^17–19^, and pointing to additional roles of hunter-gatherer mobility beyond subsistence. For example, long-distance travel for trade, resource exchange, and social ceremonies is common among extant hunter-gatherers. The Hxaro gift-giving system among the Ju/’hoansi of Botswana and Namibia involves extensive networks over distances of up to 200 km^20,21^. Archaeological^22–24^, genetic^25–27^, paleoanthropological ^28^ and paleo-environmental^29,30^ evidence also points to long-range, continental-wide interactions between hunter-gatherer groups throughout history, promoting cultural and genetic exchange.

To date, empirical studies on hunter-gatherer mobility have often focused on specific behaviors in individual camps and mostly over short time scales^31–33^. For example, Vashro and Cashdan^34^ documented gender differences in annual mobility among the Twe of Namibia and found that men travelled greater distances, possibly due to mating competition. More recently, Wood and colleagues^33^ focused on daily mobility among the Hadza and attributed larger male spatial ranges to a sexual division of labour, with men procuring rarer and more energy-dense resources (though they did not explicitly assess the reason for individuals’ trips). However, few studies have compared the relative contributions of subsistence, mating strategies, kinship ties, and other potential drivers to hunter-gatherer mobility over the lifetime. Whilst some (e.g. ^33^) have extrapolated daily mobility to make estimates of the distance travelled by individuals in their lifetimes, without direct quantification, it is impossible to determine how variability in individual-level daily behaviours scale up to regional and ultimately continental-level dynamics.

A notable exception was the pioneering work by Hewlett et al^35^ among two groups of Aka foragers in the Central African Republic: the first from Bangandu primarily subsisting from hunting and gathering, and the second from Ndelé also heavily reliant on hunting and gathering but more market-integrated and living alongside farmers within a single village. Hewlett and colleagues quantified lifetime mobility through individuals’ half ranges, or the median distance between an individual’s current residence and all places they had visited at least once in their lifetimes. In Ndelé, men’s half ranges were significantly larger than women’s, but no data on either age or gender were presented for

Bangandu. In both regions, travel was primarily driven by visiting family, hunting and gathering, or attending rituals (in addition to wage labour in Ndelé). Besides wage labour, the longest travelled distances were associated with visiting family members, supporting the hypothesis that the maintenance of social ties may determine the upper limits of spatial ranges.

Unfortunately, Hewlett et al.^35^ could not examine the effects of gender or age on mobility, possibly due to their small sample size (18 men and 24 women in Ndelé, and 41 men in Bangandu). All Central African hunter-gatherer (CAHG) populations have been generally described as patrilocal societies^36–38^, which implies that adult women should live further from their natal families (given dispersal after marriage) and that they might need to travel further to maintain family connections. However, this claim seems to be contradicted by the evidence that extant hunter-gatherers are characterized by multilocal sociality and high residential mobility of the whole family unit^2,9^. Therefore, including gender as a factor in analyses may not only clarify the reasons for hunter-gatherer mobility, but might also allow a reassessment of the claim that the BaYaka are patrilocally structured. Besides access to kin members, other potential drivers of gender differences in hunter-gatherer spatial behaviour are summarised in Table 1 (but see ^39^ for details). Testing the various hypotheses against the same dataset would provide an opportunity to establish the relative contribution of distinct factors to gender differences in mobility across the lifetime, and how their effects may vary across the geographically extensive hunter-gatherer social networks.

**Table 1.**
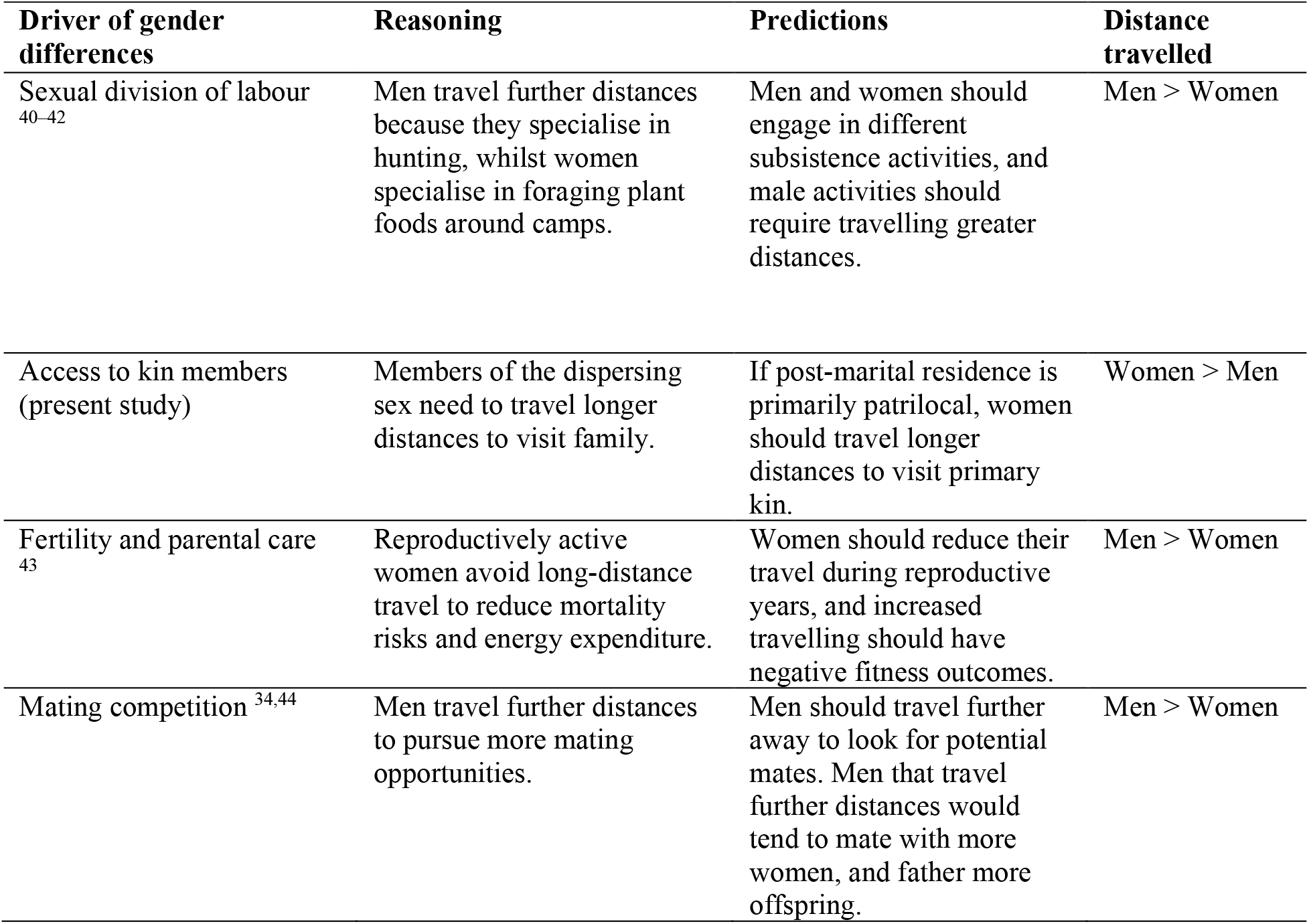
Summary of some of the main hypotheses proposed to explain gender differences in spatial behaviour.

| Driver of gender differences | Reasoning | Predictions | Distance travelled |
| --- | --- | --- | --- |
| Sexual division of labour<sup>40–42</sup> | Men travel further distances because they specialise in hunting, whilst women specialise in foraging plant foods around camps. | Men and women should engage in different subsistence activities, and male activities should require travelling greater distances. | Men > Women |
| Access to kin members (present study) | Members of the dispersing sex need to travel longer distances to visit family. | If post-marital residence is primarily patrilocal, women should travel longer distances to visit primary kin. | Women > Men |
| Fertility and parental care<sup>43</sup> | Reproductively active women avoid long-distance travel to reduce mortality risks and energy expenditure. | Women should reduce their travel during reproductive years, and increased travelling should have negative fitness outcomes. | Men > Women |
| Mating competition<sup>34,44</sup> | Men travel further distances to pursue more mating opportunities. | Men should travel further away to look for potential mates. Men that travel further distances would tend to mate with more women, and father more offspring. | Men > Women |

In the following, we present an unprecedentedly large dataset of lifetime mobility and demographic information from N=776 Mbendjele BaYaka hunter-gatherers (329 men and 443 women) across five regions of the northern Republic of Congo (Figure S1-S4). We examine gender differences in lifetime spatial ranges and how they relate to factors such as subsistence activities, kin availability and mating strategies, and investigate a possible association between gender differences in mobility and reproductive outcomes.

## Results

### Gender differences in lifetime mobility start in adolescence

We first calculated the half range of each individual^35^ and the maximum distance travelled in their lifetime. We then applied Bayesian multilevel models with a gamma distribution and log link function^33^, including random effects of residence camp nested within regions, and random slopes of gender, age and their interaction for each region to predict individual half ranges (see Materials and Methods for model specification; Tables S1-S4). Results revealed gender differences in spatial ranges over the lifetime. Women’s half ranges remained stable from adolescence onwards (Figure 1; mean posterior estimate for adolescents = 18.54km; 90% HDPI: [11.24, 25.95]; adults = 17.30km; 90% HDPI: [9.84, 24.81]). Men’s half ranges, while only moderately larger than in women during adolescence (18.51km [13.69, 24.71]), increased drastically in adults (25.83km [21.24, 30.88]; Figure 1). There was pronounced variation between regions, both in individual half ranges and gender differences (Figure S5). A similar pattern was obtained for predictors of the maximum distance during individual lifetimes (Figure S6), consistent with the rare and distant trips undertaken by men in addition to day-to-day activities (Figure S7).

**Figure 1.**
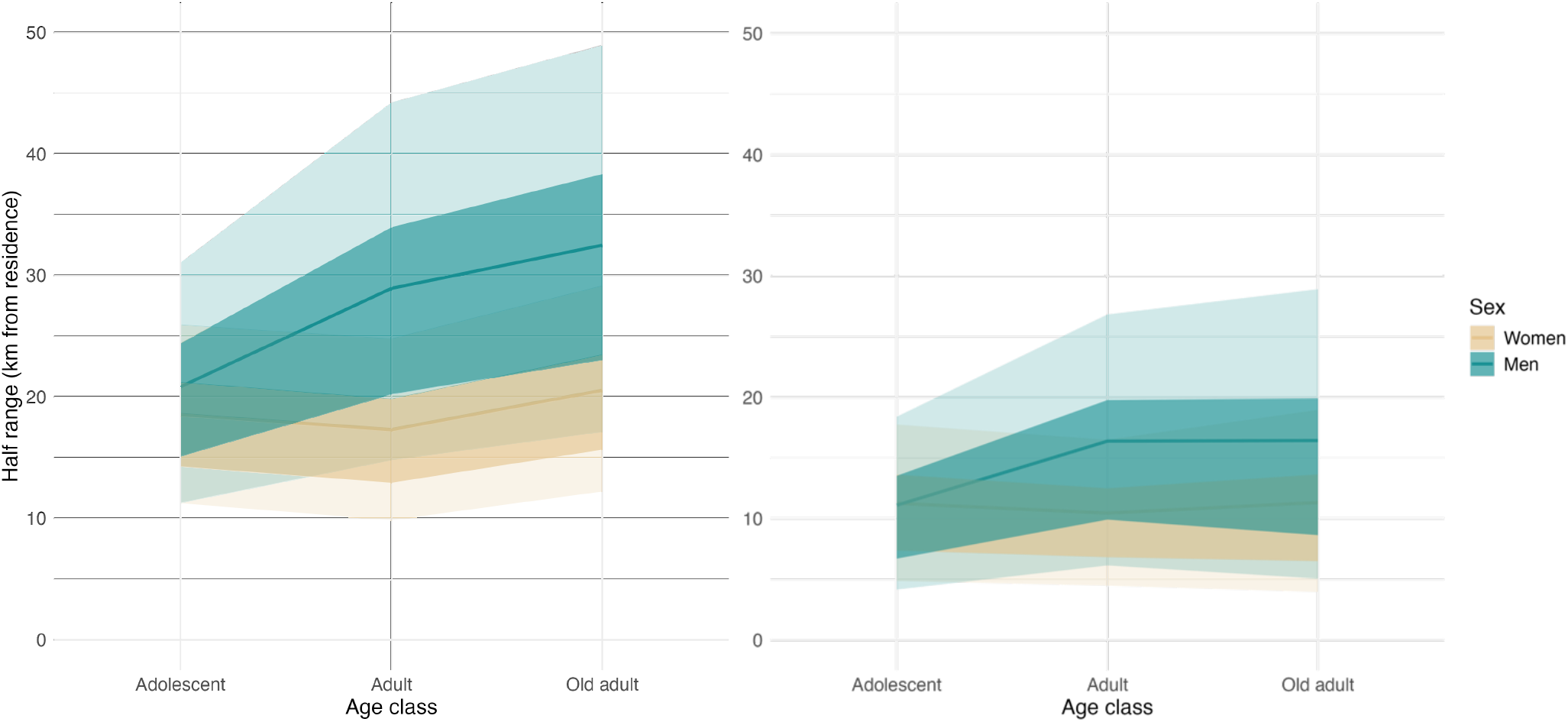
Posterior predictive distribution from Bayesian multilevel model predicting the half range of individuals as a function of age, gender and their interaction, as well as random intercepts for each residence camp nested within each region, and random slopes for both age and sex for each region. Dark line indicates posterior means, darker shaded region the 50% HPDI using the posterior standard deviation across regions and residence camps, and lighter shaded region the 90% HPDI using the posterior standard deviation across regions and residence camps. On the left, models considering all displacement and, on the right, models considering exclusively displacements walking or by pirogue and excluding those for wage labour purposes.

To obtain results more representative of selection pressures pertaining the foraging niche, we repeated the analyses after excluding trips carried out for wage labour, economic transactions and other market related activities, and only considering trips made by foot or pirogue (henceforth “traditional” displacements; cf.^34^). Men’s half ranges still increased between adolescence and adulthood (adolescents=11.11km, [4.16, 18.40]; adults=16.39km, [6.14, 26.83]). Women’s half ranges, whilst almost identical to male values in adolescence, did not increase with age (Figure 1; adolescents=11.26km [4.85, 17.66]: adults=10.46 [4.47, 16.51]). The increase in the half range of men between adolescence and adulthood was less pronounced (Figure 1; Figure S8), followed by either a plateau or slight decrease at old age. Maximum distances travelled over the lifetime showed similar values to the observed when considering all means of transport (Figure S9). Overall, our results reveal that Mbendjele BaYaka men tend to travel further distances during their lifetime than women, despite a pronounced regional variability both gender differences and individual spatial ranges highlighting the behavioural flexibility of hunter-gatherers (Figure S5-S9).

### Sexual division of labour and exploratory travel drive gender differences in mobility behaviour

We next examined potential drivers of gender differences in spatial ranges and their variability across regions with two sets of models. First we estimated the probability of individuals visiting a location due to a reported purpose of exploration, fishing, gathering, hunting, living, participating in rituals, visiting family, visiting friends, or wage labour (the most commonly cited reasons for travel), using Bayesian multilevel models with a logistic link function (Table S5). Second, we estimated drivers of differences in the distance travelled in displacements carried out for the reported purposes (Table S6).

When considering all displacements, we found that a main driver of sex differences in the probability of travelling in adults was wage labour, which in men implied both higher probability of travelling (men=0.11 [0.07, 0.17]; women=0.06 [0.04, 0.09]), and longer travelling distances (men=101.96km [59.84, 145.63]; women=42.67km [26.28, 59.87](Figure S10)). As a result, the region with the highest prevalence of travel for wage labour (Macao) also showed the largest half ranges, and largest sex differences in mobility behaviour (Figure S11-S13). The mean posterior estimate of half range in Macao was 61.69km [53.62, 79.23] for men and 29.40km [25.60, 33.44] for women, almost the same calculated by Hewlett et al.^35^ among the Aka from the N’Delé region (58.3km and 32.4km respectively). By contrast, in Minganga where travel for wage labour was rarer, the half ranges were 22.78km [18.61, 27.24] for men and 16.44km [13.61, 19.52], not far from the 27.6km obtained by Hewlett and collaborators^35,45^ among the Aka men of Bangandou (a region with no travel for wage labour, and no data on women).

When considering exclusively traditional displacements, the most common reasons for travel during adolescence were similar for both men and women, namely changing residence, participating in rituals, and visiting family and friends (Figure 2; Figure S13, S14). Although most individuals reported meeting partners during rituals, none explicitly stated that they travelled specifically to search for mates (see Figure S15 for the proportion of partners met during regular foraging activities). In adulthood, travelling for visiting family and for subsistence reasons became more prevalent for both genders.

**Figure 2.**
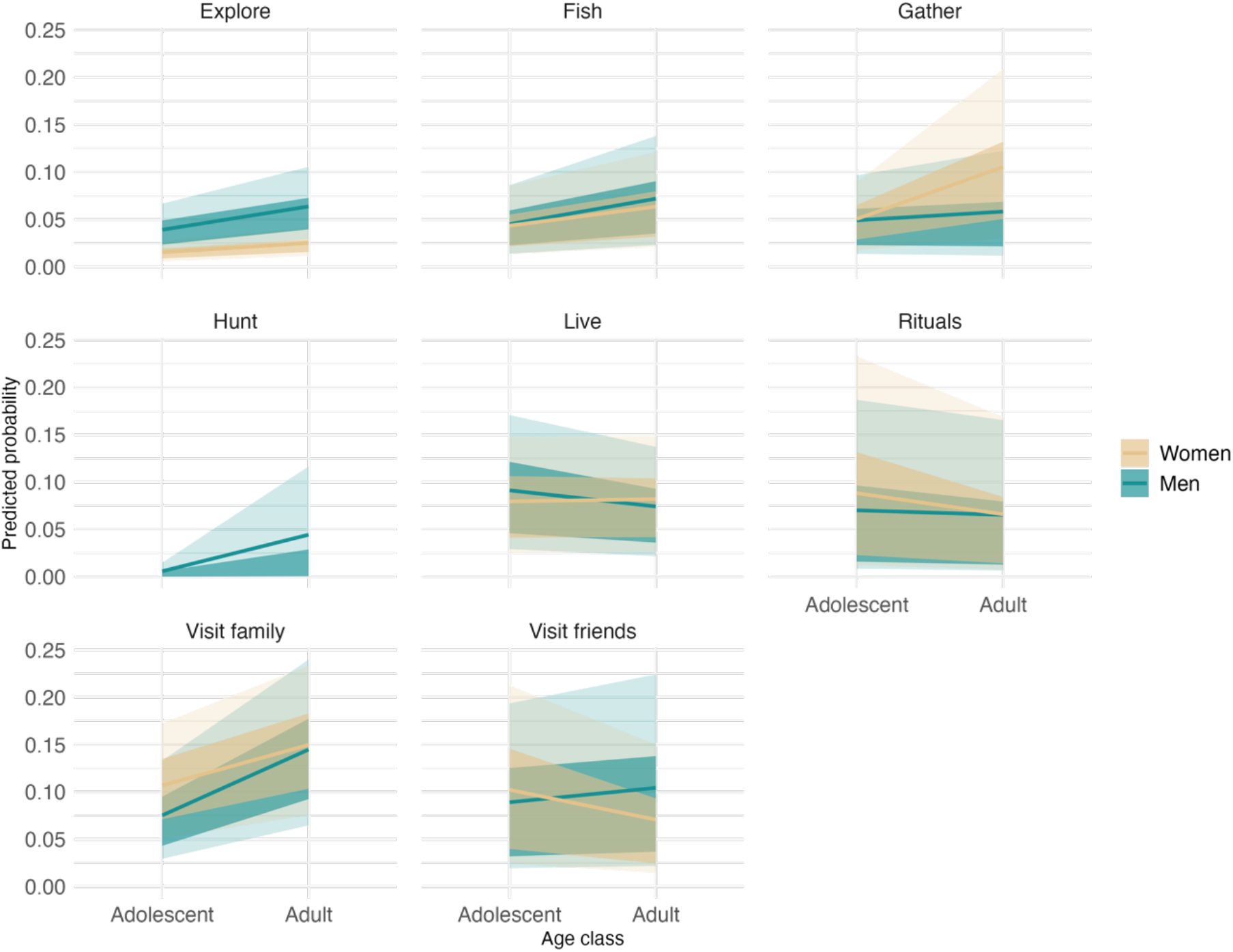
Posterior predictive distribution from Bayesian multilevel models predicting probabilities of travelling for different purposes by men and women, considering only traditional displacements. Lines indicate posterior means, darker shaded regions the 50% HDPI using the posterior standard deviation across regions and residence camps, and lighter shaded regions the 90% HDPI using the posterior standard deviation across regions and residence camps.

Women increased their travelling for gathering and fishing, whilst men did for hunting and fishing, in addition to exploratory travel.

Similar to findings by MacDonald and Hewlett^44^, we found that the average distances travelled for a given purpose were generally similar for women and men (Figure 3) and across age groups (Figure S16). This suggests that sexual division of labour (mostly in subsistence activities) may explain the increased half ranges in adult men. For example, the mean distance travelled by men to hunt (21.15km [7.28, 35.51]), a frequent and exclusive driver of male travelling, was more than double the average distance for gathering trips reported by women (9.87km [5.86, 13.93]).

**Figure 3.**
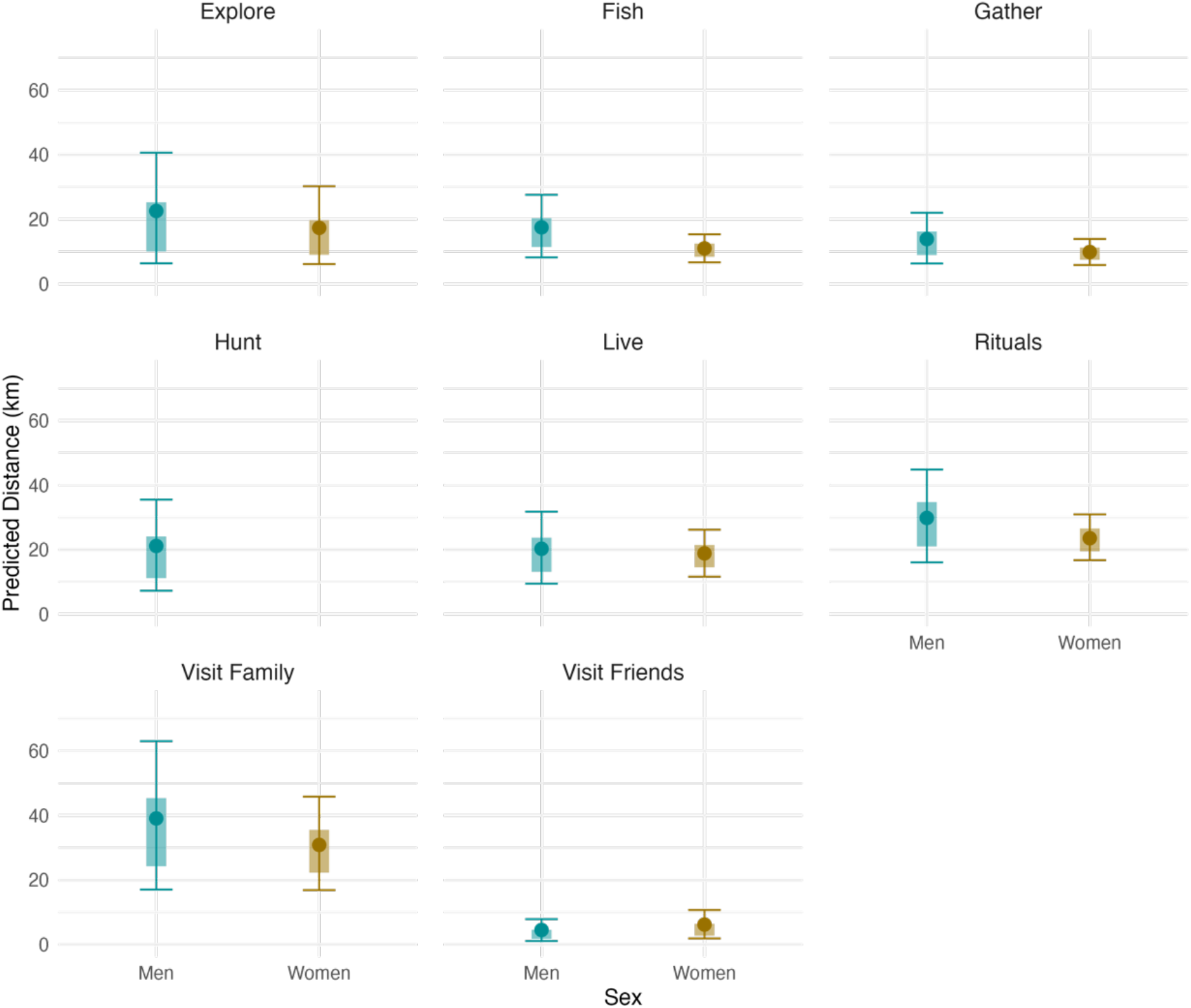
Posterior predictive distribution from Bayesian multilevel models predicting distances travelled for different purposes by men and women, considering only traditional displacements. Dark dots indicate posterior means, shaded region the 50% HDPI using the posterior standard deviation across regions and residence camps, and error bars the 90% HDPI using the posterior standard deviation across regions and residence camps.

### Men live further away than women from their primary kin

We noticed that an exception to the pattern of similar distances by age and sex for the same mobility driver is that men travelled further on average to visit kin (men=39.01km [17.04, 63.03]; women=30.87km [16.87, 45.84]; Figure 3). We therefore assessed the proximity of men and women to their close kin in two regions (Macao and Minganga) with individual data on complete fertility, birthplace, and birthplace and residence of their primary kin (N=357 individuals). We fitted Bayesian multilevel hurdle models that estimate separately the probability of zero (living with mother or father in the same camp), and probabilities of non-zero distances from camps of current residence of their father and mother. We included effects of age group and gender, as well as random intercepts for residence camp nested within regions, and random slopes for age and gender within regions (including an interaction between age group and gender did not improve model fit; see Table S7-S9 for model comparisons).

Overall, women were more likely than men to live in the same camp as their mothers (men=0.43 [0.33, 0.54]; women=0.56 [0.45, 0.66]; Figure 4). When not living with their mothers, women were on average living closer to them (men=53.11km [27.27, 84.18]; women =30.98 [15.72, 48.73]). Both men and women across age groups were less likely to live with their fathers than with their mothers (Figure 4). This might be due to higher mortality rate and lower life expectancy of men in hunter-gatherer groups including the Aka (Derkx et al. 2024; Cavalli-Sforza, 1986), or to the prevalence of sequential monogamy, with 25% (29/115) of women versus 39% (40/103) of men having been married more than once due to divorce or the death of a spouse, and to a lesser extent due to polygyny, with 21.4% (22/103) men having ever been married to two women simultaneously during their lifetimes. Finally, men did not seem to live noticeably closer to their birthplaces than women, (Figure S17-S20), and in one of the regions the opposite pattern was found. Overall, the results indicate that men seem to travel longer distances to visit their primary kin, and challenge the claims of patrilocality in the Mbendjele BaYaka.

**Figure 4.**
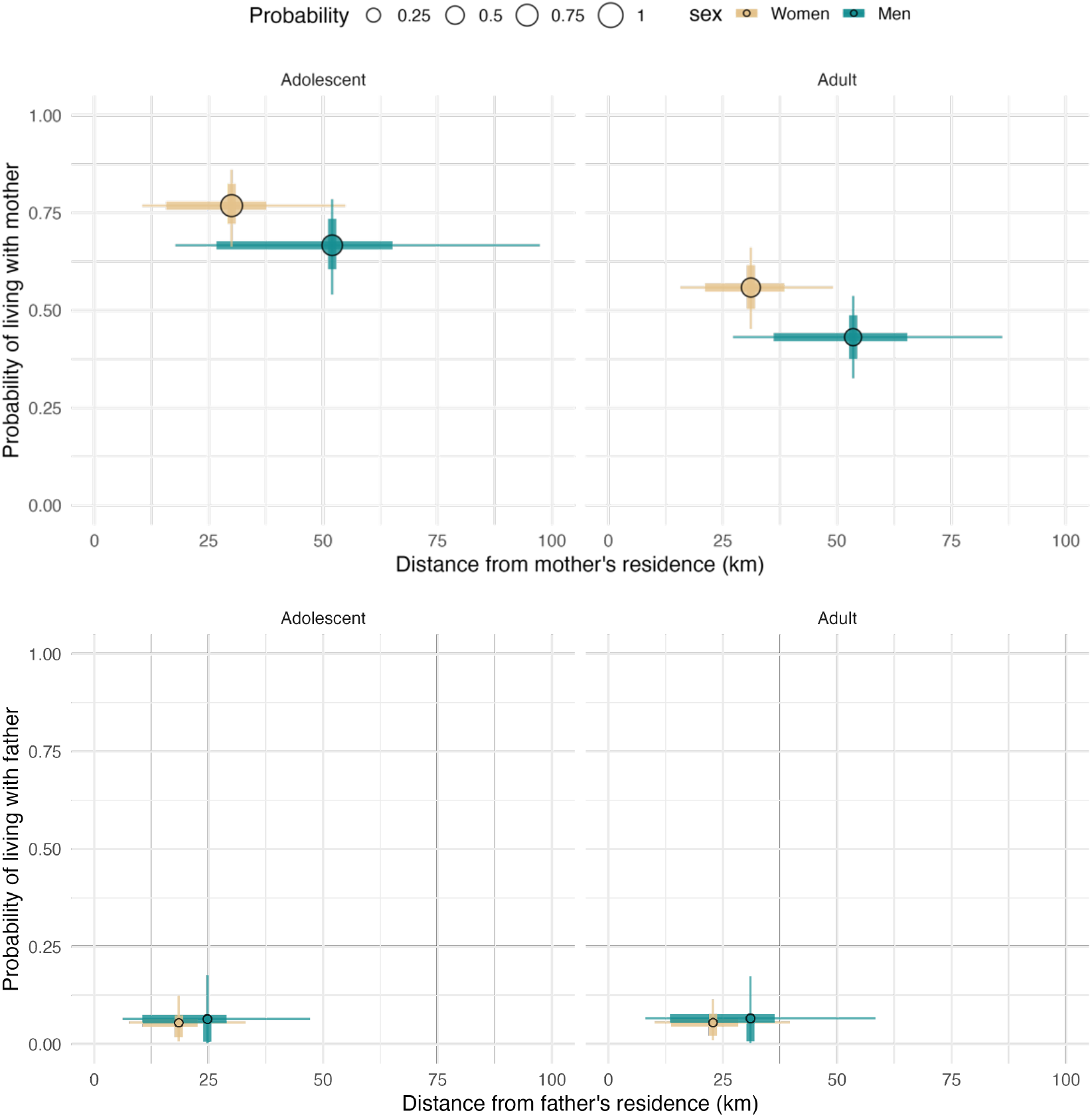
Posterior predictive distribution from Bayesian multilevel hurdle models predicting simultaneously probabilities of living in the same camp mothers (top panel) and fathers (bottom panel) by men and women as well as distances between men and women’s residence camp and that of their mothers (top panel) and fathers (bottom panel). Dot size is proportional to the probability of living in the same camp as mothers or fathers (respectively) and indicates posterior means. Rectangles indicate the 50% HDPI using the posterior standard deviation across regions and residence camps, and lighter shaded regions the 90% HDPI using the posterior standard deviation across regions and residence camps.

### Range size is associated with lifetime reproductive success in women and men

While the four discussed hypotheses imply differential mobility between women and men, two of them also explicitly predict differences at the individual level in reproductive success as a function of mobility. While the mating competition hypothesis predicts that reproductive success should be higher in the most mobile men (but not women), the fertility and parental care hypothesis implies that reproductive success should be higher in the least mobile women (but not men). To investigate the predictions, we assessed the relationship between half range and maximum distance travelled and lifetime reproductive success among the N=33 post-reproductive individuals. The restricted sample was required to isolate the effect of lifetime mobility on completed fertility and control for potential confounding effects of age^46,47,48^. Bayesian multilevel models with a Poisson link function, including random intercepts for regions and residence camp nested within regions, revealed a positive association between lifetime mobility and reproductive success for both men and women (Figure 5; Figure S21, S22), both for all data or exclusively traditional displacements (Figure S23-S26). This provides evidence against the fertility and parental care hypothesis predicting selection against women’s travel^34,49^.

**Figure 5.**
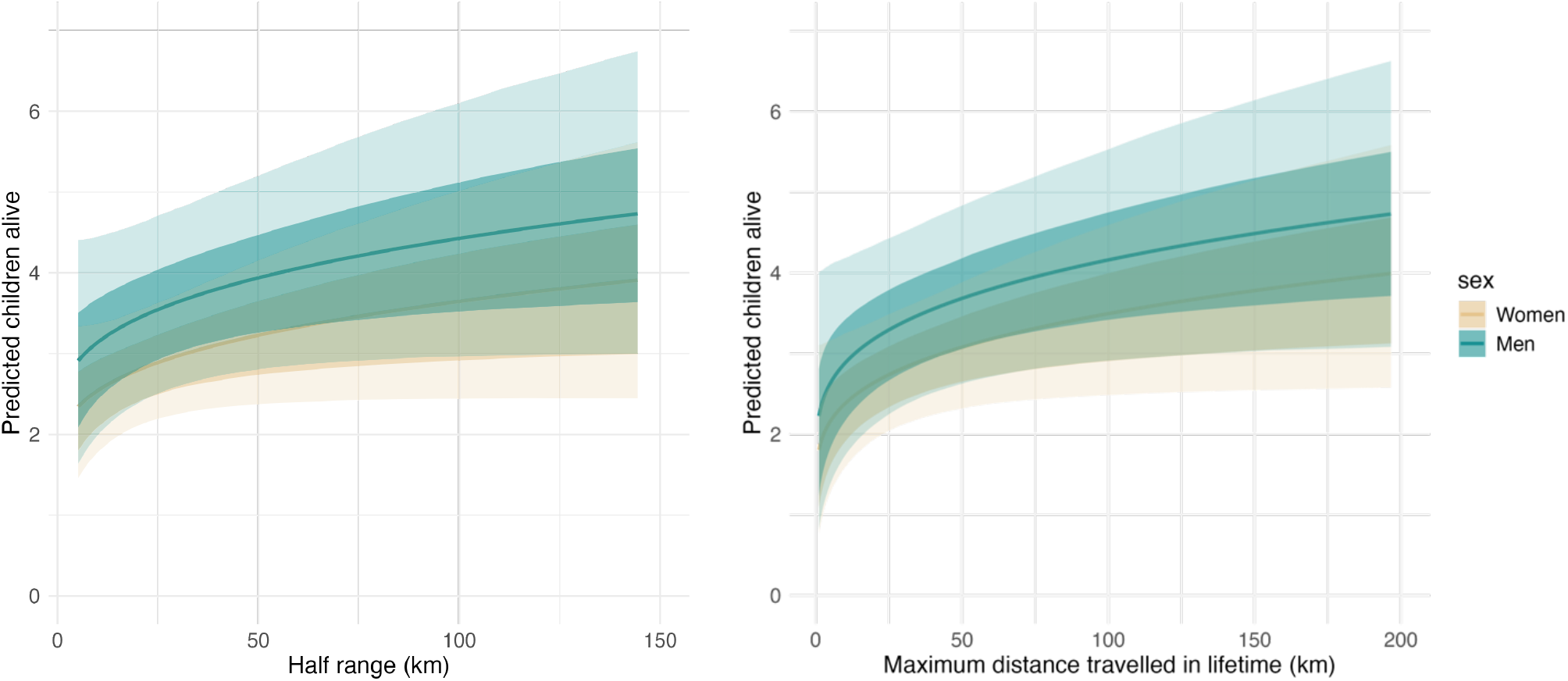
Posterior predictive distribution from Bayesian multilevel model predicting completed fertility of post-reproductive individuals as a function of sex and their half range (left) or the maximum distance travelled in their lifetime (right) considering only traditional displacements. Dark line indicates posterior means, darker shaded region the 50% HDPI using the posterior standard deviation across regions and residence camps, and lighter shaded region the 90% HDPI using the posterior standard deviation across regions and residence camps.

The results also seem to contradict the mating competition hypothesis^50,51^, which predicts an effect of mobility on fitness exclusively in men pursuing additional reproductive opportunities. Further evidence against a role of mating competition is that reproductive variance was not significantly higher in men (men=5.325; women=4.311; F=1.234, *p*= 0.219), and in general, both sexes displayed low reproductive variance compared to other hunter-gatherer societies, such as the Aka of the CAR, the Hadza, Ache, or !Kung^52^. Male-to-female reproductive skew (1.23) was also low. Although more research is required to determine the reason for the relationship between mobility and positive fitness outcomes, the similar association in both sexes suggests at least that such relationship is not the product of increased mating opportunities for men.

## Discussion

We explored drivers of gender differences in lifetime mobility among a large sample of Mbendjele BaYaka hunter-gatherers from five regions in the northern Republic of Congo. Our findings revealed noticeable gender disparities in spatial ranges beginning in adolescence and increasing in adulthood. Specifically, while women’s spatial ranges remained stable from adolescence onwards, they increased substantially in the case of men (Table 1; Figure 1). This pattern supports earlier observations by Hewlett et al.^35^ among the Aka from two regions in the Central African Republic (CAR), who noted that exploratory behaviour typically occurs between adolescence and early adulthood and stabilises thereafter. However, we showed that this is observed only in Mbendjele BaYaka men, mirroring gender differences in daily mobility previously reported only among the Hadza^33^. Gender differences in spatial ranges were reduced when only displacements representative of the foraging niche were considered, suggesting that market integration might have enhanced gender differences in mobility behaviour among the BaYaka.

More generally, our results indicate that sexual division of labour in subsistence activities mostly accounts for the larger spatial ranges observed in men compared to women^44^. BaYaka women and men tend to travel similar distances to engage in the same activity (with values similar to those reported decades ago by Hewlett et al.^35^, although they did not distinguish between means of transport), but men tend to engage in activities associated with longer distances. Furthermore, a sexual division of labour becomes more pronounced from adolescence to adulthood in the BaYaka. While adolescent men and women exhibited similar spatial ranges, mainly due to common social activities such as participating in dances or rituals (key for socializing and meeting potential partners), adult mobility was increasingly tied to subsistence particularly for men, whose exploratory behavior extended their ranges. Gathering forest products was one exception, with men traveling further on average, likely due to gender-specific specializations (for example, men gathering honey, women gathering firewood^53^). It is important to note that long-distance travel among the Mbendjele is not a new phenomenon. For example, Jacques Lalouel in 1950^54^ reported that among the Mbendjele BaYaka of Ikelemba it was not only commonplace but expected of young people to carry out extended journeys (or *mooηgơ abole*) of up to 800km during their youth.

Our results challenge previous assumptions of patrilocality and kin availability as mobility drivers among Central African hunter-gatherers^36,37^. We did not find evidence that men live closer to their birthplaces than women, and in one of the regions the opposite was true. Similarly, whilst adolescents in general tended to cohabit with their mothers, in adulthood women were both more likely to keep cohabiting with or live closer to their mothers. Men tended to travel further to visit family particularly when considering energetically costly methods such walking or pirogue travel, which might be partially explained by their mothers living further away. Therefore, although a few prior studies on kin availability and hunter-gatherer social organization^2,9,55^ have focused on patterns of camp cohabitation, our study highlights that distance to family members is also an important factor when considering kin availability.

We also found evidence contradicting the fertility-and-parental-care hypothesis^49,56^, since women who traveled more did not exhibit negative fitness outcomes, concurring with a previous study of the Twe of Namibia where nursing women travelled more to seek help from alloparents^52^. The association in both sexes between larger spatial ranges and reproductive success possibly means that more mobile individuals gain access to a greater pool of social partners and resources, rather than increase mating opportunities in the case of for men (as predicted by the mating competition hypothesis), and is consistent with the high cultural value placed on mobility in hunter-gatherer societies^57^. For example, Turnbull^58^ describes how the Mbuti refer to lacking relatives in other bands as “walking emptily”. Moreover, during visits to other camps, one becomes a full member of the band and acquires the same rights to work and hunt there, which may further value mobility and possibly reinforce it. Alternatively, having more children might also lead to increased travel needs, such as more frequent hunting and gathering trips or seeking help from alloparents.

In summary, our study provides novel evidence of the interaction between social and subsistence factors driving hunter-gatherer mobility. The extent of the variability in mobility patterns across individuals, age groups, genders or regions, highlights that flexibility in moving behaviours might be a key component to the adaptive potential of the foraging niche^20,59,60^. Future work is required to quantify precise responses in individual mobility strategies to changing ecological, social or economic environments^61^. Similarly, additional research is also needed to connect our findings on the lifetime spatial ranges of individuals with the shorter, daily and yearly movements that also constitute a large part of hunter-gatherer mobility. These include trips to procure raw materials, food, firewood, or to gather social information from nearby camps or seasonal relocations^20,33,61^. These displacements likely have significant evolutionary implications and contribute to the overall structure of hunter-gatherer mobility. Such integration of mobility data at different scales is required to better understand the selection pressures that have shaped hunter-gatherer mobility over evolutionary time and to explore the broader implications for human biological and cultural evolution.

## Materials and methods

### Study population and data collection

The Mbendjele BaYaka are a hunter-gatherer population living in the rainforests of the Congo Basin, with an estimated population of 15–20,000^38^. They belong to a broader group of hunter-gatherers collectively known as Central African hunter-gatherers ^62,63^ or rainforest hunter-gatherers ^64,65^ that have inhabited the rainforests of West and Central Africa since at least the last interglacial ^29,66^.

For the present study, we surveyed five different regions in the Sangha and Likouala districts of the Republic of Congo. All regions were located within forest environments and around 150km apart from one another. Hunter-gatherer populations from all regions are highly residentially mobile, forage daily for most of their subsistence needs, speak the same language, rely on the same resources, and seasonally work in agriculture for neighbouring farming populations. Data collection was conducted between February 2022 and June 2023. All adolescent and adult individuals within the five study regions were surveyed, comprising a sample size of 776 participants across 113 camps. Camp sizes across regions were close to the mean of 19.2 individuals observed in a recent global survey of 249 ethnographic hunter-gatherers^15,67^(Figure S2) and were comprised on average of 4.65 huts (min=1, max=25), inhabited by on average 4.70 people (min=1, max=25).

Data were collected using a structured questionnaire that captured demographic information (such as gender, age category, place of birth, time spent in current residence, languages spoken, and exposure to Bantu populations), mobility history, reproductive data, and kin availability. The questionnaire was administered in Lingala (the second language for most participants) and French by the first, second, and third authors. In cases where a participant was not fluent in Lingala, a Mbendjele field assistant facilitated translation to ensure the clarity and accuracy of responses. In the Macao and Minganga regions, participants (N=357) were additionally asked about reproductive and marriage practices, including completed fertility, providing a deeper context for examining reproductive success in relation to mobility.

### Mobility data collection and distance calculations

To document lifetime mobility, participants were asked to indicate whether they had visited a predetermined list of locations—temporary camps, villages, and regions at varying distances from their current residence—based on a prior survey of locations common in Mbendjele mobility (Following the framework developed by Hewlett et al.^35^ and later ^34^). For each location, participants specified the reason (or reasons) for travel, the mode of transport, and the number of times visited. The great-circle distance between each pair of locations (in kilometers) was calculated using the Haversine formula on the Earth’s surface, acknowledging this approach may underestimate actual travel distances, as Mbendjele paths through the dense rainforest are often sinous, particularly when on foot or by canoe^32^.

### Ethical approval and consent

Approval for this research was obtained from the Ethics Commission (Ethikkommission) of the University of Zurich (Permit Nr. 20.2.8), the Canton of Zürich, as well as by the Presidency of Research and Cooperation from the Marien Ngouabi University, the Ministry of Justice, Human Rights and promotion of indigenous peoples of the Republic of Congo, and the Ministry of the Environment and Sustainable Development of the Republic of Congo. All participants provided informed consent after receiving a comprehensive explanation of the study’s objectives, the data to be collected, and the broader research goals. This information was presented orally, with questions answered to ensure that consent was fully informed.

### Statistical approach

We employed Bayesian inference for all statistical analyses, which allows for comprehensive interpretation of posterior distributions and provides interval estimates that reveal the range of plausible values for each parameter. For all models, we report the 50% and 90% highest density posterior intervals (HDPI), which represent the narrowest interval containing 50% and 90% of the posterior probability for each parameter ^68^.

Regularizing priors were adopted in order to prevent the model from overfitting the data ^68^, and prior predictive checks were performed to make sure all priors were consistent with characteristics of each parameter. In addition, we fit alternative parameterizations for all models, to confirm that the results presented below were qualitatively robust to changes in priors. Parameter estimation was achieved with the *brms* (v2.22.0) R package ^69^, running three Hamiltonian Monte Carlo chains in parallel until convergence was suggested by a high effective number of samples and R^ estimates of 1.00. Trace plots were also inspected to make sure chains converged to the same target distributions and posterior predictions were compared to the raw data to confirm the match between model predictions and the descriptive summaries of the samples.

For model comparisons, we used Leave-one-out Cross-validation with Pareto smoothed importance sampling to regularize importance weights implemented with the *loo* R package ^70^.

### Half ranges and distance travelled in individuals’ lifetimes

We modeled predictors of individuals’ *half-ranges* (the median distance between their place of residence and all places visited at least once^35^) using Bayesian generalized linear mixed-effects models with a Gamma distribution to account for the positive, continuous nature of the data ^33^. The model included age, sex, and their interaction as fixed effects, with random slopes for age and sex at the regional level and a nested random intercept for residence camps within regions to account for hierarchical structure and between-group variability. The model is specified as follows:

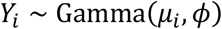

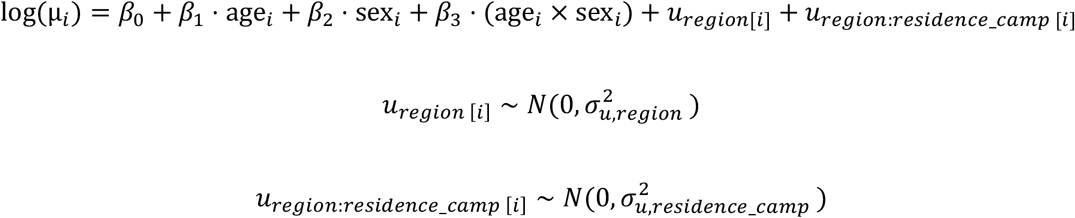

Priors (for model including only traditional displacements):

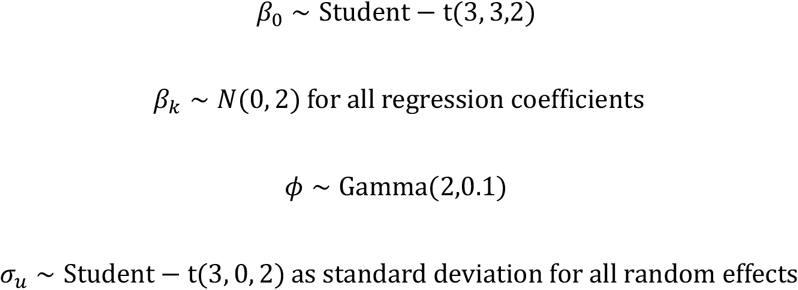

For the maximum distance travelled by individuals throughout their lifetimes, models were identical, with the exception htat the prior for the intercept was:

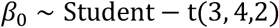

Models considering all displacements can be found in the accompanying GitHub repository.

### Probability of travelling for specific activities

We modeled the probability of travelling for specific activities (fishing, hunting, visiting family, visiting friends, participating in rituals, gathering, living or exploring) as a proportion of the total places visited, using Bayesian hierarchical binomial models. Each response variable was defined as the number of times a particular activity was cited as the reason for visiting a place, with the total number of places visited as trials. These models account for the hierarchical structure of the data by including random intercepts for both region and residence camp nested within regions and random slopes for sex and age at the regional level to capture regional variability in the effects of these predictors.

The full model formula is:

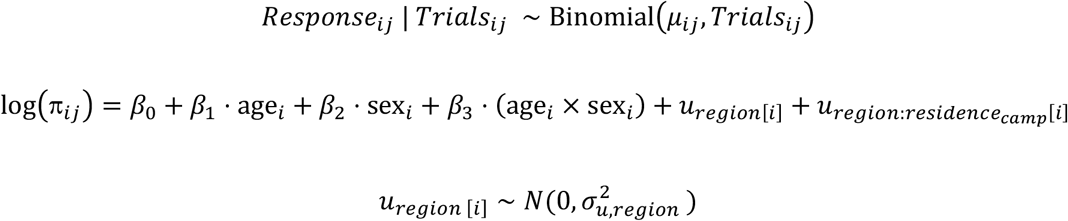

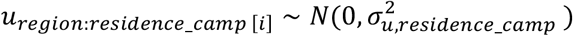

Priors (for model including only traditional displacements):

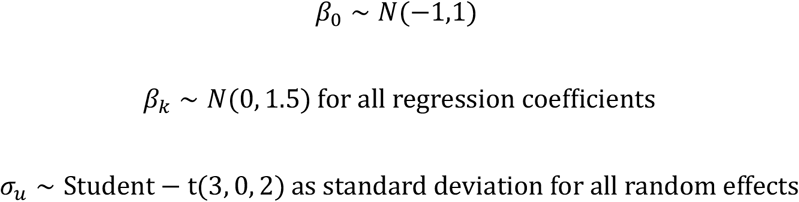

To assess the robustness of our findings, we fitted three model types one including random intercepts for region and residence camps, one including an interaction between age and sex, with random slopes for sex and age at the regional level, and random intercepts for residence camps nested within regions; and a third one keeping the same random slope structure but excluding the interaction. Models including the interaction were selected as the best models (Table S5) Models considering all displacements can be found in the accompanying GitHub repository.

### Distance travelled for different activities

We modeled the differences in distance traveled across all displacements for which at least one purpose from the eight most popular ones (fishing, hunting, visiting family, visiting friends, participating in rituals, gathering, living or exploring) was mentioned (N=1459 displacements) by fitting Bayesian generalized linear mixed-effects models with a Gamma distribution, suitable for the positive and continuous nature of the data. The model included age, sex, and activity type as fixed effects, along with random slopes for sex and activity at the regional level. Additionally, nested random intercepts were specified for residence camps within regions and a unique reference identifier to account for hierarchical structure and individual-level variability. Additional models including an interaction between sex and activity were fitted but model comparisons did not provide support for the interaction effect (Table S6). The best model is specified as follows:

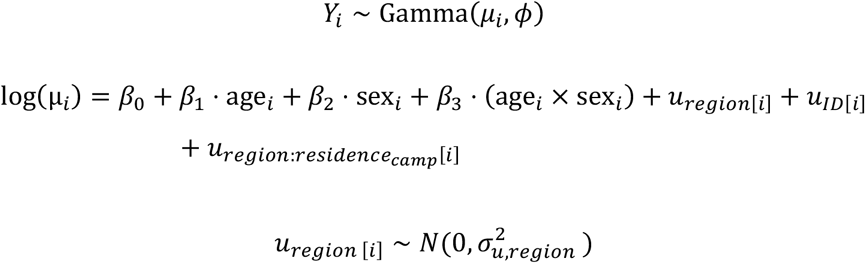

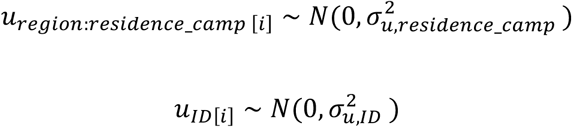

Priors (for model including only traditional displacements):

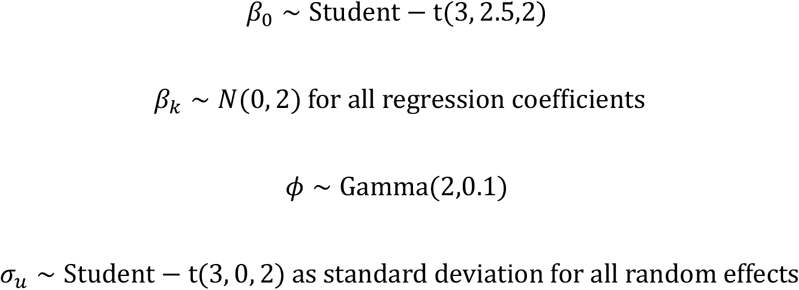

Models considering all displacements can be found in the accompanying GitHub repository.

### Distance from kin

To investigate the factors influencing the distance individuals lived from their mother’s residence, their father’s residence and their own birthplaces, we employed Bayesian hurdle gamma models. These models are well-suited for data with a large number of zeroes (i.e., individuals living with their mother/ father/ at their birthplace) and positively skewed continuous distances for those who live further away^71^. We included age group and sex as fixed effects, random intercepts for both region and residence camp nested within regions and random slopes for sex and age at the regional level. We also fitted similar models with an interaction between age and sex but including the interaction did not improve model fit (see Table S7-S9 for model comparisons). The model formulas for the best models were as follows:

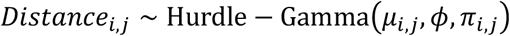

Where *μ*_*i,j*_ is the mean of the possitive distance (gamma component), *ϕ* is the shape parameter of the gamma distribution, and *π*_*i,j*_ is the probability of zero distance (zero component).

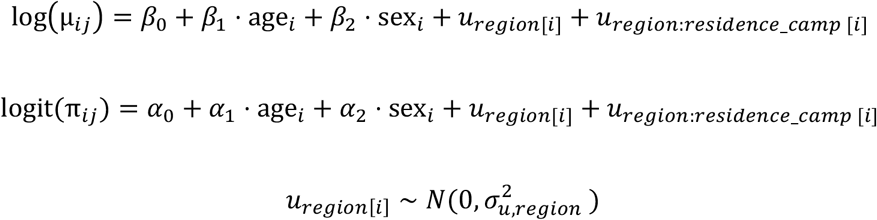

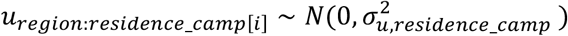

Priors (for models predicting distance from maternal and paternal residence):

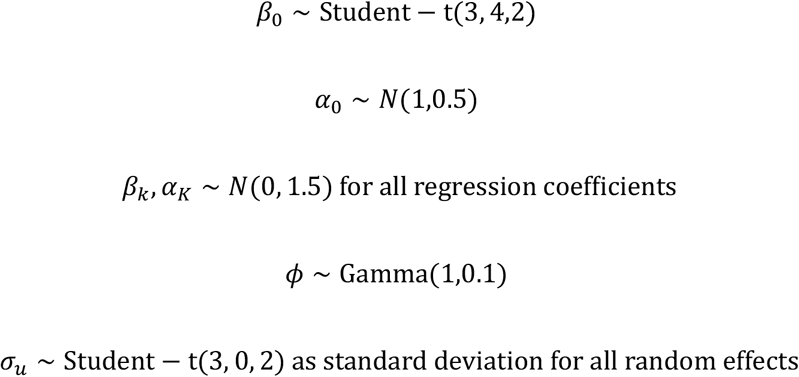

Priors (for models predicting distance to own birthplace):

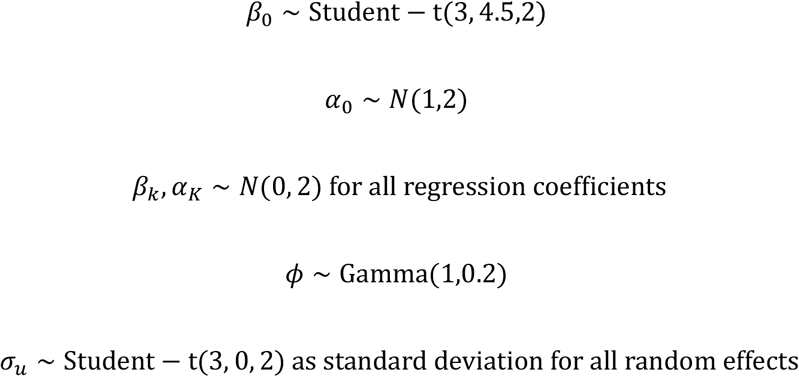

### Relationship between travel and reproductive success

To examine the relationship between mobility and reproductive success, we modeled the number of living offspring (i.e. completed fertility) for post-reproductive individuals as a function of lifetime mobility, represented by the standardized log-transformed half range and sex. To account for hierarchical data structure, we included random intercepts for regions and residence camps nested within regions (we did not include random slopes due to the reduced number of post-reproductive individuals). For models including all individuals (and not only post-reproductives), we also included age as predictor in our model. The model was specified as follows:

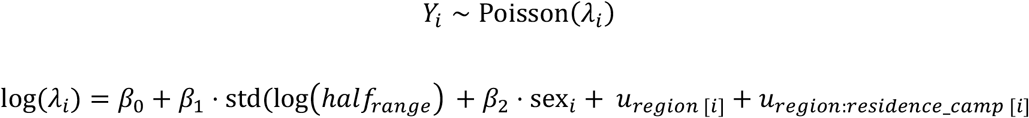

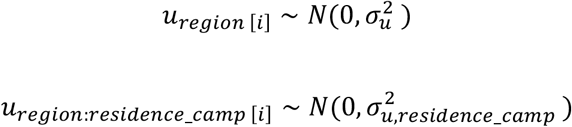

Priors:

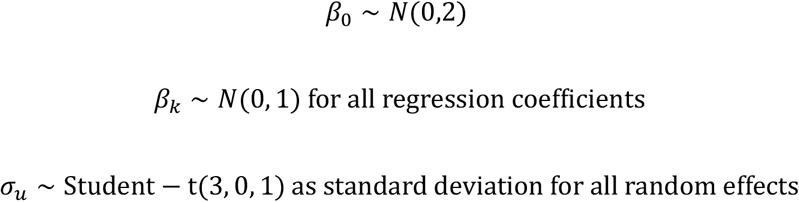

Models including the maximum distance travelled by individuals in their lifetime as predictor (instead of half range) were similar, and the predictor was also log-transformed and standardised. Models considering all displacements can be found in the accompanying GitHub repository.

## Acknowledgements

The authors wish to acknowledge all the BaYaka who generously agreed to participate in the study, the village chiefs of Macao and Minganga for facilitating our stay and Rudy Schlaepfer for help with fieldwork preparation. CP-I also wishes to acknowledge the Fundación La Caixa (grant LCF/BQ/EU19/11710043), A.H. Schulz Foundation and Leakey Foundation for funding the project. CP-I, LV and ABM are greateful to the University of Zürich for funding this project.

